# The *in vitro toxicity* of nitrile and epithionitrile derivatives of glucosinolates from rutabaga in human and bovine liver cells

**DOI:** 10.1101/419960

**Authors:** Ian Latimer, Mark Collett, Zoe Matthews, Brian Tapper, Belinda Cridge

## Abstract

Previous evidence suggests that select nitrile and epithionitrile derivatives of glucosinolates can cause liver disease in cows grazing on brassica forage crops. A toxic incidence in New Zealand in cattle grazing brassica led us to investigate the direct *in vitro* hepatotoxicity and possible inhibition of the ABCG2 transporter of five nitrile compounds. In this study, we investigated 1-cyano-2-hydroxy-3-butene (CHB, epithionitrile derivative of progoitrin), 1-cyano-2-hydroxy-3,4-epithiobutane (CHEB, nitrile derivative of progoitrin), 3-butenenitrile (nitrile from sinigrin), 4-pentenenitrile (nitrile from gluconapin), and 5-hexenenitrile (nitrile from glucobrassicanapin). Cell viability was assessed following 24- and 72-hr treatments with the 5 different compounds using the MTT assay (HepG2 cells and bovine primary liver cells). Additionally, ABCG2 transporter function was assessed. The results showed that none of the tested compounds caused cytotoxicity at concentrations up to 2 mM for 24hr. Over 72-hr the maximum concentration was 20 μM but no reduction in cell viability was observed. No inhibition of the ABCG2 transporter occured at concentrations up to 1 mM. Overall this study suggests that direct or secondary toxicity due to selected nitrile or epithionitrile derivatives of these glucosinolates was not the cause of the toxic event in cattle.

## 1 Introduction

### Background

In the spring of 2014 in the Southland and South Otago regions of New Zealand there was a large and unprecedented outbreak of sudden deaths, photosensitization, reduced body condition, increased incidence of metabolic disease, and reproductive problems in dairy cattle grazing rutabaga (*Brassica napus* ssp. *napobrassica*, swedes)(1). These crops were virtually weed-free and the rutabaga plants were well-grown, leafy with long stems, and were starting to flower. Daily access by cattle to the crops was restricted (break feeding) according to time or calculated consumption per animal. This was an unusual poisoning scenario as NZ cattle routinely feed on this crop with no observed toxicity. Photosensitization was the most outstanding clinical presentation of many of the cows and was secondary to liver disease as indicated by elevated serum liver enzyme activities. Histopathological lesions of the liver in many of the cows that died or that were euthanized showed distinctive but subtle lesions in small interlobular bile ducts, variable portal fibrosis and bile duct hyperplasia, as well as mild fatty change or patchy necrosis in the parenchyma (2). These lesions closely resembled those seen in bulb turnip (*B. rapa*) photosensitization (3). Kidney lesions were variable and comprised tubular dilation, cast formation, and scattered tubules showing epithelial necrosis (2). Following the 2014 outbreak, an epidemiological investigation was carried out by DairyNZ. Samples of flower, leaf, stem and bulb were analysed for 21 different glucosinolates known to be found in swedes, turnips and rape crops. The concentration of one glucosinolate, progoitrin (25 μmol/g dry matter), was 10-50 times higher than any of the other 20 (<2 μmol/g dry matter) (1).

### Brassicaceae and glucosinolates

The Brassicaceae family of plants contain a wide variety of glucosinolate secondary compounds. Over 130 glucosinolates have been identified, and each brassica species or cultivar characteristically contains a variety of different glucosinolates that vary in their proportion and concentration in different plant parts (e.g. leaves, stems and flowers) and under different growing conditions (4). When brassica plant cell walls are disrupted by chewing, the endogenous myrosinase enzyme converts the glucosinolate compounds into a range of metabolites. The metabolites produced are dependent on both the parent glucosinolate(s) and the conditions of the environment in which the degradation takes place. In addition, the pH of the medium where this degradation occurs is important; breakdown at a low pH (<4) produces predominantly nitrile metabolites and at higher pH isothiocyanates (5). This complexity means that it has been difficult to determine the full spectrum of biological actions of the glucosinolate metabolites (6).

It has been hypothesized that nitrile and/or epithionitrile derivatives of glucosinolate compounds from turnip (*B. rapa*), and rape (*B. napus* ssp. *biennis*) forage crops cause hepatotoxicity or cholangiotoxicity in cattle (7). When crambe (*Crambe abyssinica*) and rapeseed meals were fed to rats, bile duct and liver and renal tubular epithelial cell damage resulted and this was attributed to the nitriles (8). The same lesions were found in rats fed the epithionitrile CHEB from *epi*-progoitrin (9). Crambe seed meals have been reported as having high concentrations of the parent glucosinolate *epi*-progoitrin; which formed the nitrile 1-cyano-2-hydroxy-3-butene (CHB) and two diastereomeric isomers of the epithionitrile, 1-cyano-2-hydroxy-3,4-epithiobutane (CHEB) at low pH (8). Rapeseed (*B. napus*) meal reportedly have high concentrations of progoitrin and hydrolysis produced the same daughter compounds as crambe seed but with the (*R*) configuration (8).

This evidence led us to investigate the *in vitro* toxicity of the progoitrin-derived nitrile (CHB) and epithionitrile (CHEB). Since the nitriles produced by sinigrin (the dominant glucosinolate in Brussels sprouts, cabbage and kale), gluconapin and glucobrassicanapin (3-butenenitrile [3-B] or allyl cyanide, 4-pentenenitrile [4-P] and 5-hexenenitrile [5-H], respectively) were readily available, we chose to investigate them as well. The synonyms, formulae, and structures of the investigated compounds and their parent glucosinolates are shown in Table 1.

**Table 1.**
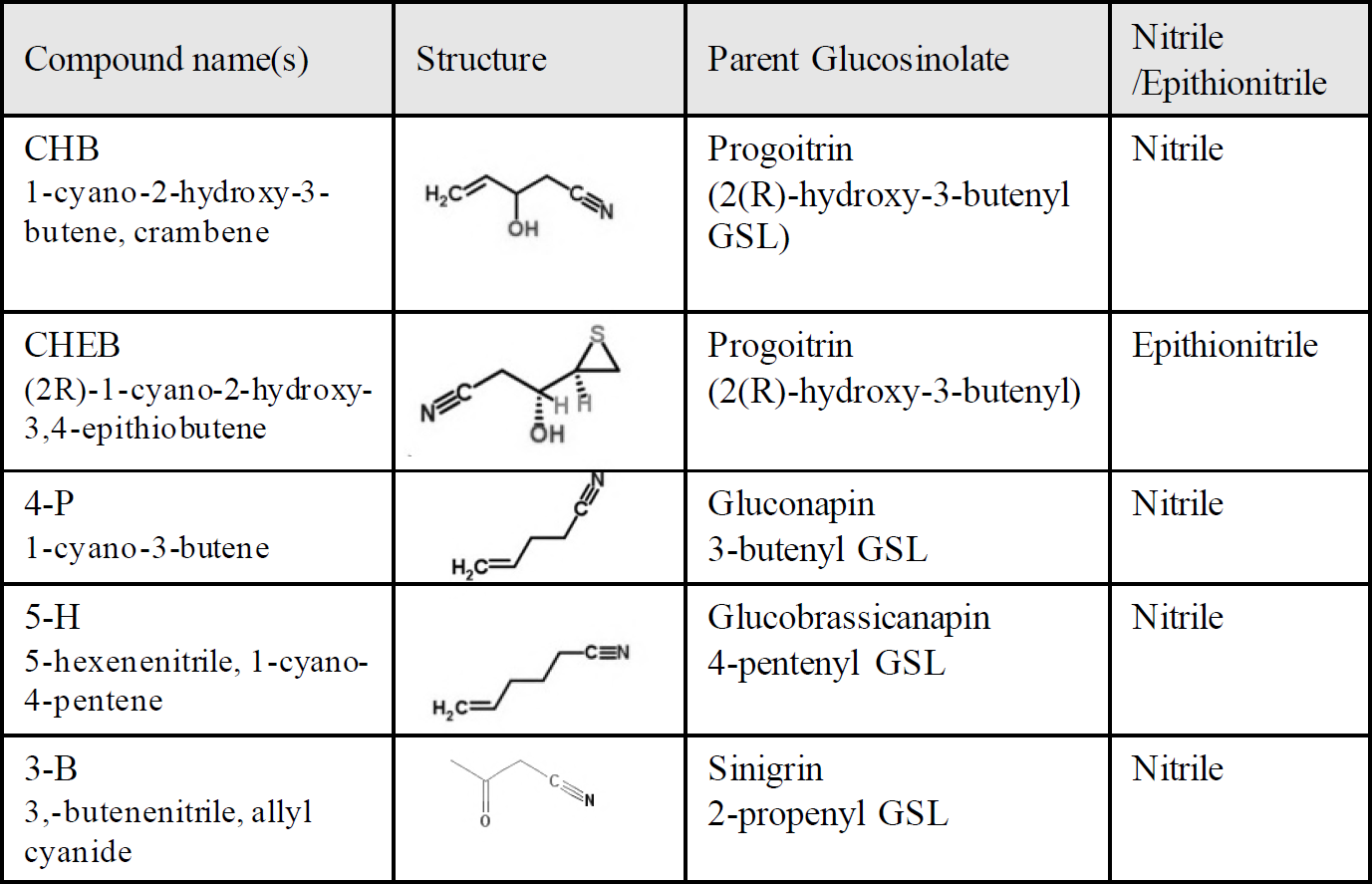
Compounds used in the current study with their parent glucosinolate and representative structural form.

### Photosensitization

Clinical reports from the poisoning incidence highlighted a significant degree of photosensitization in poisoned animals (2). This was determined to be a secondary feature of the observed liver damage. Certain types of liver damage, especially when bile ducts are involved, leads to the disruption in the normal biliary excretory pathway of dietary chlorophyll breakdown pigments. These pigments include pheophorbide *a* and phytoporphyrin (phylloerythrin) (10). In the liver, the adenosine triphosphate (ATP)-binding cassette (ABC) transporter G2 (ABCG2), also known as the breast cancer resistance protein (BCRP), actively transports both of these pigments into the bile (11, 12). This implies a potential role for ABCG2 in the pathogenesis of the phototoxicity. We therefore hypothesized that one or more of these nitriles could inhibit ABCG2 and therefore prevent the normal biliary excretion of chlorophyll derivatives leading to photosensitization. This aspect of the liver metabolism of nitriles has not been previously investigated.

## Materials and Methods

### Materials

The two progoitrin derivatives, 1-cyano-2-hydroxy-3-butene (CHB) and (2R)-1-cyano-2-hydroxy-3,4-epithiobutane (CHEB), were custom synthesized by BDG Synthesis (Wellington, New Zealand) and certified to at least 97% purity. 3-butenenitrile (3-B), 4-pentenenitrile (4-P) and 5-hexenenitrile (5-H) were purchased from Sigma-Aldrich, NZ. HepG2 cells were a generous gift from Greish laboratory group, Department of Pharmacology and Toxicology, University of Otago, Dunedin, New Zealand. Bovine liver samples were sourced from Silver Fern Farms Ltd NZ, Balclutha, New Zealand. Advanced Dulbecco’s Modified Eagle’s Medium (DMEM), Fetal Bovine Serum (FBS), GlutaMAX-I, Hanks’ Balanced Salt Solution (HBSS), Hepes-HBSS Buffered Salt Solution (pH 7.8) containing 1.78 mM NaHCO_3_, 5.5 mM D-Glucose, 10 mM HEPES, ATB solution containing Penicillin G (Na-Salt) 300,000 IU, Gentamycin 0.15 g, Streptomycin 0.20 g, PBS 300 mL were sourced from GIBCO^TM^, ThermoFisher (NZ). (3-[4,5-Dimethylthiazol-2-yl]-2.5-Diphenyltetrazolium Bromide (MTT), Sterile Water, Crystalline Bovine Insulin (zinc), Collagenase II (390 units/mg), Trypan Blue, Trypsin (0.25%), Fumitremorgin C (FTC), Dimethyl Sulfoxide (DMSO), Phosphate Buffered Saline (PBS), Ethylene Glycol-bis(β- aminoethyl ether)-N,N,N’,N’-Tetra-acetic Acid (EGTA) were purchased from Sigma Aldrich (NZ). The ABCG2 vesicles and fluorescent substrates kit were purchased from GenoMembrane (Yokohama, Kanagawa, Japan). Pall 96-well glass filter plates 1.0μm, glass fibre were sourced from AcroPrepTM, Pall Corp. EGTA-PBS contained EGTA 1 mM in PBS adjusted to pH 7.0 before filter sterilization.

### Cell culture and cytotoxicity assay

HepG2 hepatocellular carcinoma cells were cultured in Advanced Dulbecco’s Modified Eagle Medium (DMEM) supplemented with 1% FBS and 20 mL GlutaMAX-I supplement. Cells were seeded into 96 well plates at 8,000 cells/well for cytotoxicity and allowed to adhere for 24-hr. Solutions were prepared from the pure compounds as follows. CHB was diluted in sterile water to a 2 mM stock before dilution directly in cell media. CHEB, 3-B, 4-P, and 5-H were prepared in DMSO solutions (2 mM stock) and added to media to make a diluted final solution. All compounds were prepared for final in-well concentrations of 1, 2.5, 5, 10, 15, 20, 100, 200 μM, and 2 mM. Treatments were added in volumes of 10 μl to the plate across six wells each following adhesion and incubated for 24 or 72 hr. Untreated control was 10 μl of media with 1 % solvent (DMSO or sterile water). The MTT assay (13, 14), was performed following 24-hr/72-hr incubation. The combination MTT assay involved adding 20 μM of each compound in combination. All experiments were performed in triplicate with four experimental repeats.

### Bovine liver primary cells protocol

The protocol was a revised version of previously published protocols (21). Liver samples were minced with trauma shears and washed with 20 ml antimicrobial ATB solution. Tissue was homogenized in 50 ml HBSS until no large pieces of tissue remained and the sample had the consistency of a thick slurry. 25 μl of collagenase II 1 mg/ml was added before the mixture was moved into a cell culture hood and stirred gently for 12 min. The cells were then filtered through cheesecloth and centrifuged at 1080 rpm for 5 min at 4 ^°^C. The supernatant was discarded carefully so as not to disturb the pellet and 20 ml of EGTA-PBS solution was added to the tube. The tube was centrifuged again at the same settings and time. The supernatant was discarded again and another 20 ml of ATB was added for the final spin. The remaining pellet was re-suspended in DMEM media and a sample was taken for trypan blue exclusion assay for cell viability (15). All methods were carried out in accordance with the relevant guidelines and regulations. Ethical approval was not required for these studies as the sample were sourced from a commercial abattoir, however the University of Otago Animal Ethics Committee was informed of the work and the protocols being used in this study.

The best cell culture results followed two days of incubation allowing time for viable cells to attach to the flask. Twenty-four hours after initial culture the media was discarded and fresh media was put on, cells were then left for two days before seeding for MTT. Trypsin and re-seeding of flasks were found to reduce the number of viable cells and so flasks were seeded at an initial concentration that eliminated the need for new flasks within the test period. Seeding densities were determined during resuspension in DMEM with viable cell number determined by the trypan blue assay.

### ABCG2 vesicle transporter assay

As bovine (*Bos taurus*) ABCG2 transporters were not available for the transporter assay, an alternative model had to be found. It was determined that commercial preparations of human, rat, or mouse ABCG2 were available and so genetic alignments were conducted to determine the best model. A Basic Local Alignment Search Tool (BLAST) of the Uniprot database found the human ABCG2 gene to be 84.5% identical to bABCG2, whereas of the other experimental options, mouse was 79.9% and rat was 79.6% identical. The important functional domains of ABCG2 are the nucleotide binding domain, and the integral transmembrane domain (16). A Uniprot alignment comparison between ABCG2 for bovine (*Bos taurus*) and human (*Homo sapiens*) found identical sequences at the nucleotide binding domain and only 11 differences in sequence in the transmembrane domain, with only 4 of those being significantly dissimilar. Therefore, the human ABCG2 vesicles were purchased for studying ABCG2 inhibition. The ABCG2 assay was performed following the protocol of the manufacturer. Briefly, a reaction mix containing 10μl of vesicles 9.5μl of Buffer A, 20μl of 10mM MgATP solution, 5μl of 100μM Lucifer Yellow (LFY), and 5.5μl of test compound was incubated for 5 min at 37^°^C. Following incubation, 200μl of chilled Buffer B was added to stop the reaction. Filter suction was used to remove the reaction mix and 5 washes were performed using Buffer B (200μl/well). LCY contained within the vesicles was recovered using three washes of 50μl 10% SDS centrifuged at 2000rpm for 1 min each time. Finally, 100μl DMSO was added to each well and read at excitation wavelength of 427nm and emission wavelength at 535nm on a Clariostar^TM^ plate reader. Results of treatments were compared to DMSO negative control and FTC (20μM) positive control. Compounds were individually tested at 200μM (4 wells each) and 1mM (2 wells each).

### Statistical Analysis

All statistical analysis was performed with Graph Pad Prism 6 Software. All treatment groups were compared to vehicle control using a one-way analysis of variance (ANOVA) with Dunnett’s multiple comparison post-hoc test. Where applicable, Tukey’s multiple comparison test was used to compare between all treatments including the control. All tests had significance set at p <0.05.

## Results

### 24 hour treatment of HepG2 cells

None of the five compounds showed any cytotoxicity by MTT assay (Fig. 1) in the HepG2 hepatocarcinoma cell line at concentrations of 100 μM, 200 μM, or 2 mM. When treated for 24-hr, there was no significant difference between the control solvent-only and the treatments with the compounds.

**Figure 1:**
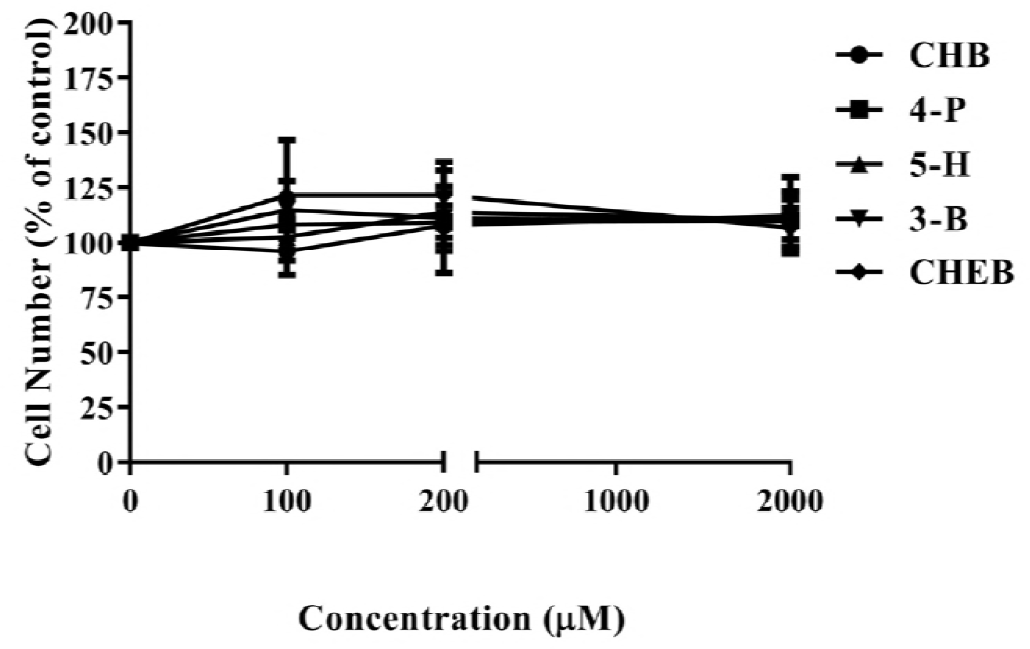
Graph of results of 24 hr MTT cytotoxicity assay of CHB, CHEB, 3-B, 4-P and 5-H in HepG2 cells. Data is cell viability expressed as a percent of solvent-only in media control (0 μM). Data expressed as Mean ±SEM with α=0.05 for significance. No test treatments showed any significant difference from control. n=3 repeats.

### 72 hour treatment of HepG2 cells

To extend the investigation, HepG2 cells were also exposed to compounds for a period of 72hr. In this model of sub-chronic exposure, doses were decreased to be more relevant to an *in vivo* situation. Under these conditions no compound showed any cytotoxicity by MTT assay (Fig. 2). There was no significant difference between the control solvent-only and the treatments with the compounds.

**Figure 2:**
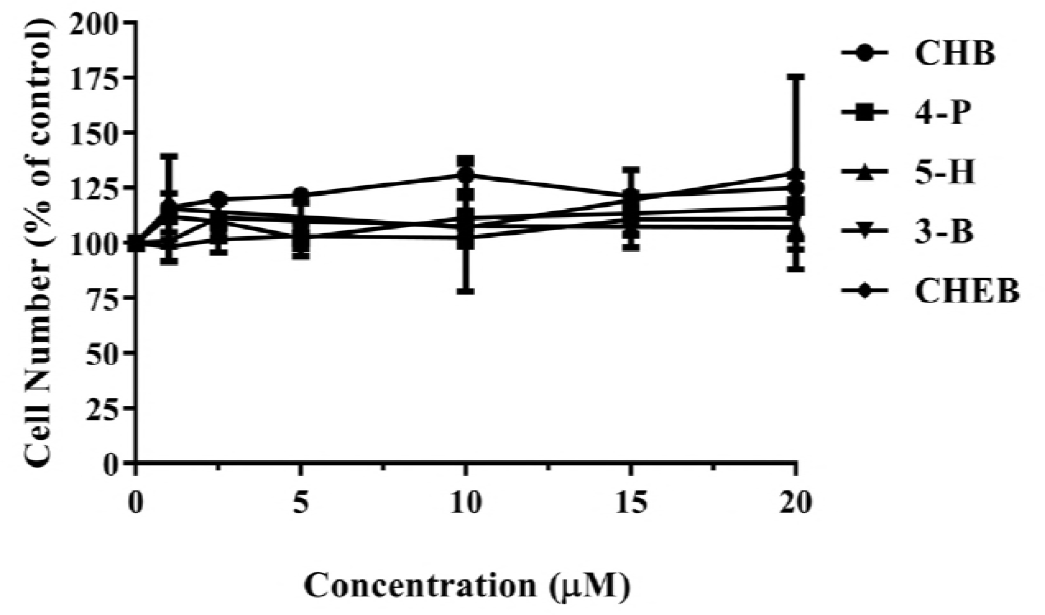
Graph of results of 72 hr MTT cytotoxicity assay of CHB, CHEB, 3-B, 4-P and 5-H in HepG2 cells. Data is cell viability expressed as a percent of solvent-only in media control (0 μM). Data expressed as Mean ±SEM with α=0.05 for significance. No test treatments showed any significant difference from control. n=3 repeats.

### Combination treatment of HepG2 cells

To test for possible synergism between the individual compounds, which would have been ingested simultaneously, a combination treatment was performed. Each chemical was cross-tested in a dual exposure (equal concentrations) with each of the other chemicals and as overall mixture containing all the test compounds. HepG2 cells were exposed to the mixtures for 72-hr at concentrations up to 20 μM. Again, no toxicity in terms of cell death was reported with any treatment (Fig. 3).

**Figure 3:**
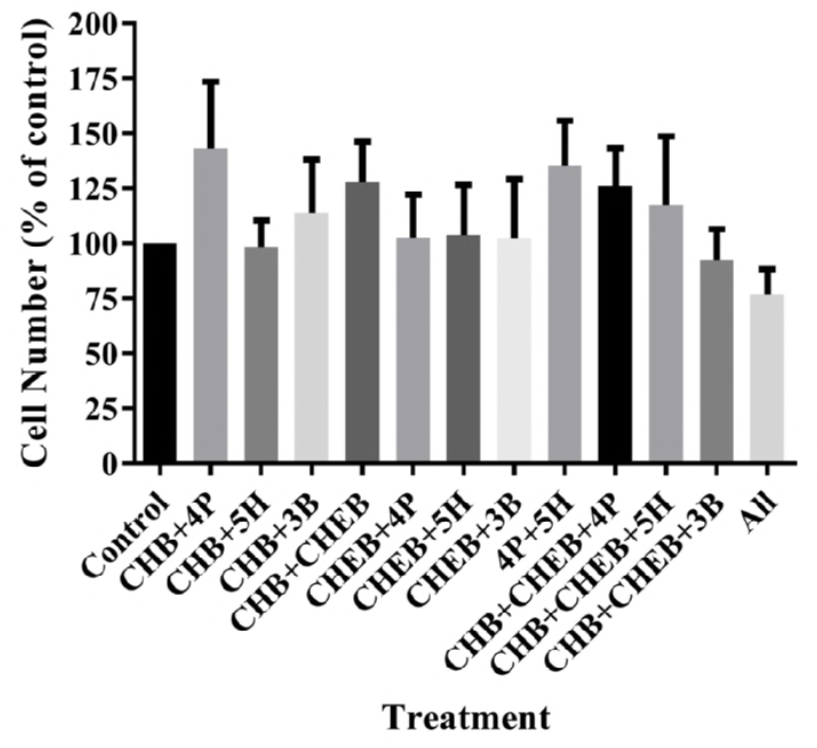
Graph of results of 72 hr MTT cytotoxicity assay of combinations of CHB, CHEB, 3-B, 4-P and 5-H. Data is cell viability expressed as a percent of solvent-only in media control (0 μM). Data expressed as Mean ±SEM with α=0.05 for significance. No test treatments showed any significant difference from control. n=4 repeats.

### Bovine liver primary cells

As the poisoning event had occurred in cattle and clinical features of bile duct damage were reported, each of the compounds was also screened in a primary cell preparation derived from cattle liver. The culture was prepared to ensure that it maintained all hepatic cell types including the bile ductal cells and the Kupffer cells. However, once again, no compound showed any cytotoxicity by MTT assay (Fig. 4) at concentrations of 100 μM, 200 μM, or 2 mM.

**Figure 4:**
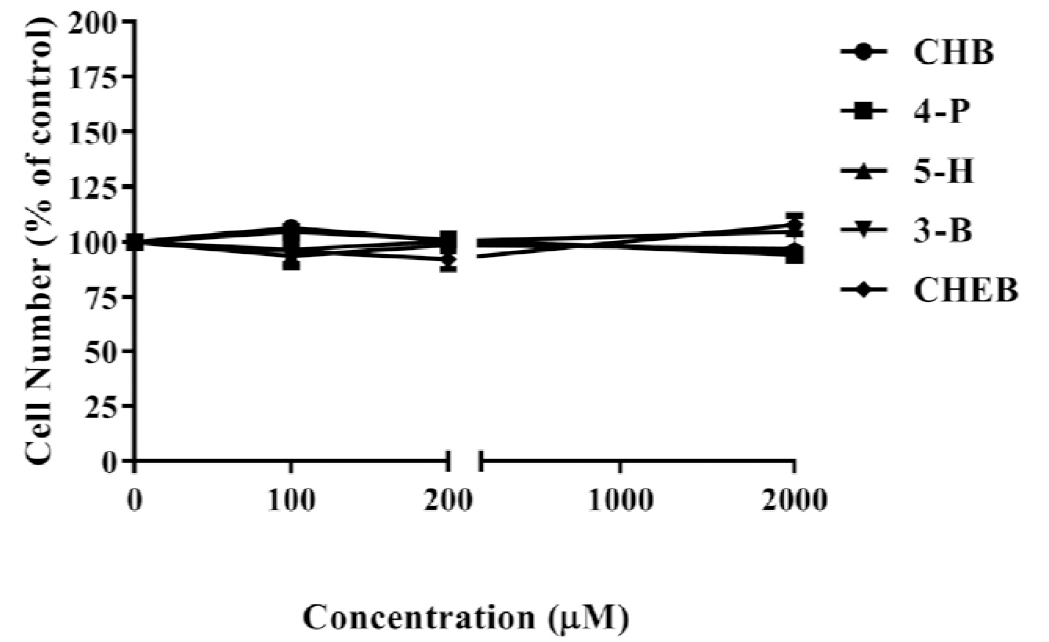
Graph of results of MTT cytotoxicity assay of CHB, CHEB, 3-B, 4-P and 5-H in primary bovine liver cells. Data is cell viability expressed as a percent of solvent-only in media control (0 μM). Data expressed as Mean ±SEM with α=0.05 for significance. No test treatments showed any significant difference from control. n=3 repeats.

### ABCG2 transporter assay

As reported phototoxicity may have been caused by inhibition of the ABCG2 transporter, the ability of each compound to specifically inhibit this transporter was tested. Each compound was incubated with the ABCG2 vesicles at concentrations of 200 μM and 1 mM for 5 minutes (optimised exposure time for this assay). None of the compounds reduced the ability of the ABCG2 transporter to move the fluorescent substrate across a vesicle membrane indicating they caused no significant inhibition in this system (Fig. 5).

**Figure 5:**
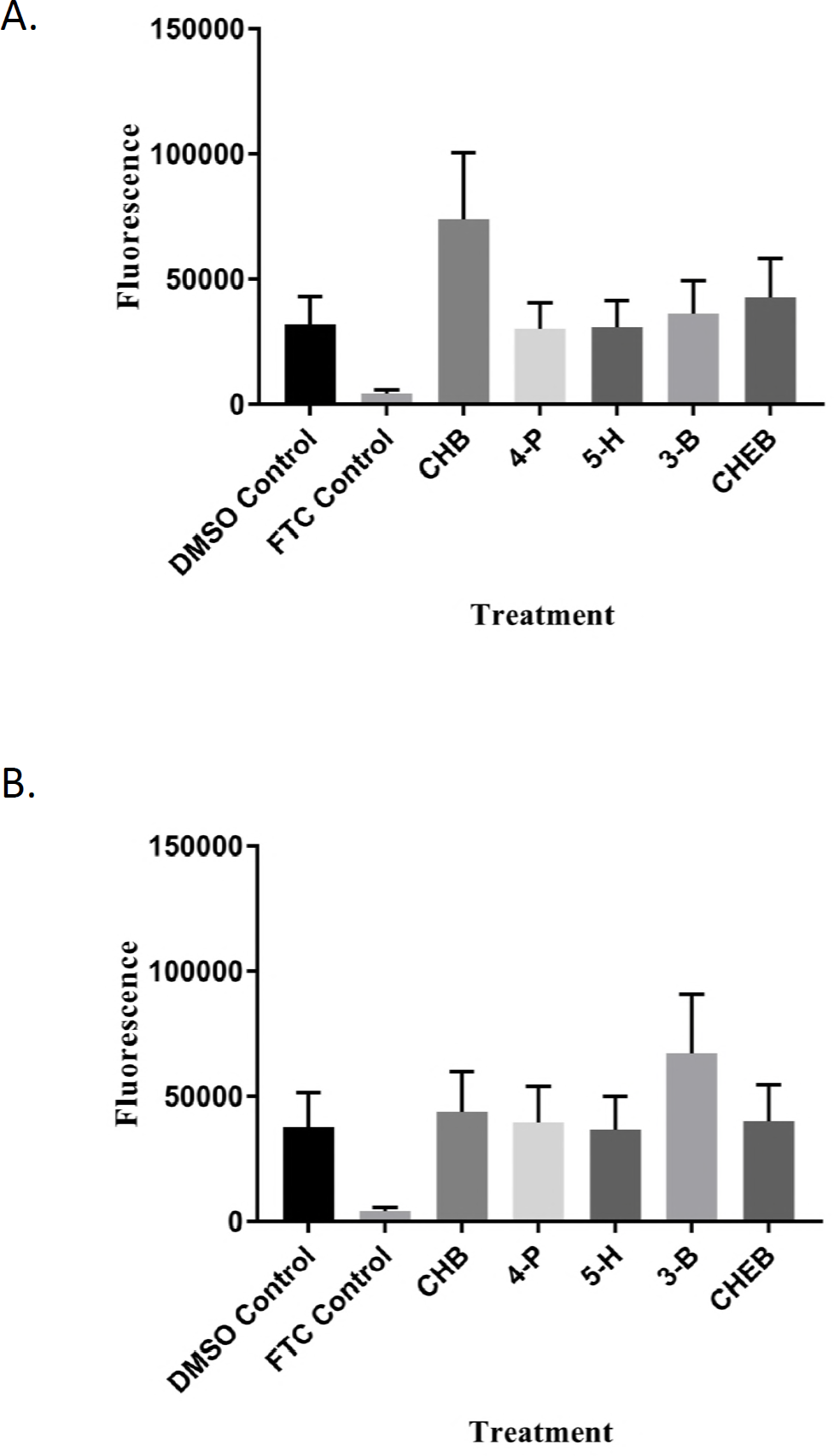
Graphs of results of ABCG2 inhibition assay at a) 200 μM and b) 1 mM. Data represented as Mean ±SEM with α=0.05 for significance. FTC is the positive control fumitremorgin c. No test treatments showed any significant difference from control. n=4 for test compounds (a). n=2 for test compounds (b)

## Discussion

### Direct toxicity

It is clear from the results that none of these nitriles or epithionitrile glucosinolate-derivatives have any direct cytotoxic effect on HepG2 cells or primary bovine liver cells *in vitro*. High IC_50_ values for 3-B (3-butenenitrile) and 4-P (1-cyano-3-butene) have been previously reported in rat liver cells (17). This study reports IC_50_ values of 510 μM and 530 μM for 3-B and 4-P respectively which is higher than we found in the HepG2 cell line but is well beyond the expected level of nitrile that would be experienced via the diet. Kelleher *et al*, also tested 1-cyano-2,3-epithiopropane (CETP) the epithionitrile derivative of sinigrin (c.f. the nitrile derivative of sinigrin, 3-B, used here) and again observed low toxicity (IC_50_ of 770 μM) in rat liver cells (17).

In contrast, 1-cyano-2,3-epithiopropane (CETP) was found to be toxic in HepG2 cells with an IC50 of 32 μM (18). However, full cytotoxicity was not reported with highest dose tested (370 μM) reducing cell viability to approximately 15% of control (18). In contrast, in primary murine liver cells concentrations of 300 μM only reduced cell number to 50% of control (18). However, it is interesting to note that the study used synthetic CEP in the murine cell test but a brassica extract in the HepG2 test. This suggests that the difference in reported toxicity may be influenced by the extraction process as well as the species. In the current study and in the work by Kelleher (2009), synthetic versions of each chemical were used which may be less toxic *in vitro*. However, none of the three *in vitro* studies to date have showed any significant toxicity, at physiologically relevant exposures, for the nitrile derivatives of sinigrin or gluconapin.

### ABCG2 transporter

It is clear from the results of the ABCG2 transporter assay that none of these glucosinolate derivatives inhibit ABCG2 at concentrations up to 1 mM. This is the first study to investigate the effects of these compounds on the ABCG2 transporter. As high levels of ABCG2 is closely correlated with multi-drug resistance in cancer cells, numerous studies have investigated the structural requirements for inhibition of this protein. The protein structure of the ABCG2-FTC complex has recently been published (19). This shows that FTC binds into the active site (competitive inhibition) and prevents conformational changes required for the transportation activity (19). It is presumed that this is the primary mode of inhibition due to the fact that the most potent ABCG2 inhibitors contain several key structural similarities which resemble the FTC molecule (20). To date, there is little evidence that the presence of a nitrile or epithionitrile species alone is predictive of ABCG2 inhibition. However, the current study did not include an evaluation of the nitrile/epithionitrile metabolites of the glucosinolate indole-3-carbinol (21) which have a structural backbone more closely resembling known ABCG2 inhibitors. The evaluation of these compounds against ABCG2 activity may be an avenue for further study.

### Conclusions

The results of this study indicate that direct liver cell toxicity or the inhibition of ABCG2 transporters in the liver by nitrile or epithionitrile derivatives of progoitrin and three other glucosinolates was not the likely cause of the cattle deaths, photosensitivity or liver disease in the poisoning outbreak in New Zealand in 2014. We have been unable to show any evidence of *in vitro* toxicity of CHB, CHEB, 3-B, 4-P, or 5-H. This suggests that toxic mechanism is something that we are unable to replicate *in vitro* or alternative metabolites are responsible for the toxicity.

## Contributions

All studies were conducted and analysed by I.L. under the direct supervision of B.C., the lead author and researcher for this paper. M.C., Z.M. and B.T. provided invaluable advice and feedback throughout the work and assisted with the revision of the manuscript.

